# Sex-Specific neurogenic and cognitive responses in a murine model of accelerated aging

**DOI:** 10.1101/2025.11.25.690416

**Authors:** Ricardo Gómez-Oliva, Andrea Chamorro-Francisco, Isabel Atienza-Navarro, Livia Carrascal, María D. Freire-Aragón, Rosario Hernández-Galán, Pedro Nunez-Abades, Mónica García Alloza, Carmen Castro

**Affiliations:** Área de Fisiología, Facultad de Medicina, Universidad de Cádiz, Cádiz, Spain; Instituto de Investigación e Innovación Biomédica de Cádiz, Cádiz, Spain; Departamento de Fisiología Facultad de Farmacia. Unicversidad de Sevilla, Sevilla, Spain; Hospital Universitario Puerta del Mar de Cádiz, Cádiz, Spain; Departamento de Química Orgánica. Universidad de Cádiz, Cádiz, Spain

**Author notes:** Both authors contributed equally to the work (Ricardo Gómez-Oliva, Andrea Chamorro-Francisco).

## Abstract

Aging is associated with cognitive deterioration accompanied by a reduction in hippocampal neurogenesis. Murine models have been widely used to study aging and age-related cognitive decline, as they recapitulate many of the key features of the degenerative process. Among these, the SAMP8 strain represents a well-established model of accelerated aging, characterized by early-onset and progressive cognitive impairment, Alzheimer’s disease-like neuropathology, and an initial increase in hippocampal neurogenesis that ultimately depletes the neural stem cell pool. Notably, most studies using murine models of aging or neurodegeneration have focused on males or mixed-sex cohorts, leaving sex-specific differences in neurogenesis and cognitive decline largely unexplored. Recent evidence indicates that diterpenoid treatment ameliorates cognitive decline and enhances neurogenesis in 6-month-old male SAMP8 mice. However, whether females exhibit similar responses remains unknown. In this study, we characterized sex differences in hippocampal neurogenesis in 6-month-old SAMP8 mice and examined potential sex-dependent effects of diterpenoid therapy. Our findings reveal marked sex differences in neurogenic capacity and treatment responsiveness. While diterpenoid treatment enhanced hippocampal neurogenesis and cognitive performance in males, these effects were largely absent in females. Overall, female SAMP8 mice exhibited reduced baseline neurogenesis and a diminished response to therapy, underscoring the importance of considering biological sex in the design of therapeutic strategies for age-related cognitive decline.

## INTRODUCTION

Aging is associated with a progressive decline in hippocampal neurogenesis (Babcock, Page, Fallon, & Webb, 2021; Gomez-Oliva, Martinez-Ortega, et al., 2023), which has been implicated in cognitive deterioration and neurodegenerative disorders such as Alzheimer’s disease (AD) (Liu et al., 2023) (Terreros-Roncal et al., 2021). Several murine models have been used so far to study the effects of aging on cognitive performance and neurodegeneration. The SAMP8 mouse model, characterized by accelerated aging and cognitive deficits, has been widely used to study age-related hippocampal dysfunction (Diaz-Moreno et al., 2018; Diaz-Moreno et al., 2013). Besides exhibiting multisystemic signs of aging, this model displays an early-onset and progressive decline in cognitive function, which becomes evident around 4 months of age (H. Yagi, Katoh, Akiguchi, & Takeda, 1988). From 6 months onward, SAMP8 mice begin to show neuropathological symptoms commonly associated with Alzheimer’s disease (Butterfield & Poon, 2005). Notably, an initial increase in neurogenic activity in the dentate gyrus of the hippocampus (DG) occurs during early stages, which ultimately leads to a depletion of the neural stem cell (NSC) pool. This exhaustion appears to be driven by extrinsic signals within the neurogenic niche (Diaz-Moreno et al., 2013; Gang et al., 2011; Soriano-Canton et al., 2015).

Recent findings suggest that cognitive resilience in aging is closely linked to the functional responsiveness of neurons generated in the DG during adulthood (Montaron, Charrier, Blin, Garcia, & Abrous, 2020). These studies indicate that memory performance in older individuals depends not only on the quantity of adult-born neurons, but also on how effectively these neurons are activated and integrated into cognitive processes (Montaron et al., 2020). However, most studies describing the relevant role of neurogenesis in neuropathological aging have been performed in male subjects (Diaz-Moreno et al., 2018; Diaz-Moreno et al., 2013; Gomez-Oliva, Geribaldi-Doldan, et al., 2023) or using subjects from both sexes without analyzing the influence of sex (Liu et al., 2023). Thus, sex differences in neurogenesis and its response to stimuli remain poorly understood, despite increasing evidence that aging and neurodegeneration differ between males and females (Lee, Cevizci, Lieblich, & Galea, 2025) (S. Yagi et al., 2020). Notably, recent studies have identified sex-specific differences in adult hippocampal neurogenesis. Males show a faster maturation and greater attrition of newly generated neurons in the DG compared to females (S. Yagi et al., 2020), suggesting that neurogenesis in males may be more sensitive to environmental and stimuli modulation (Chow, Epp, Lieblich, Barha, & Galea, 2013). These findings emphasize the importance of incorporating sex as a key biological variable in experimental research on neuroplasticity and cognitive aging.

Understanding the impact of sex on DG plasticity is essential for developing effective therapeutic strategies; however, sex-specific responses to therapeutic interventions are understudied aspects of neurobiology (Hyer, Phillips, & Neigh, 2018). Many preclinical studies fail to include both sexes, yet pharmacological and lifestyle interventions often yield different outcomes in males and females (Knufinke, MacArthur, Ewald, & Mitchell, 2023; Waters, Network, & Laitner, 2021). This is particularly relevant in aging research, where therapies aimed at enhancing neurogenesis or slowing cognitive decline may be differentially effective depending on biological sex. Identifying these differences could lead toward sex-specific treatments adapted to the individual aging trajectories of male and female individuals.

A recent report shows that diterpenoid therapy ameliorates cognitive decline in 6-month-old male SAMP8 mice by enhancing hippocampal neurogenesis (Gomez-Oliva, Martinez-Ortega, et al., 2023). However, it is uncertain whether such treatment produces similar effects in females. In this study, we aimed to characterize sex differences in hippocampal neurogenesis and cognitive performance in 6-month-old SAMP8 mice and to determine whether diterpenoid treatment responses differ between sexes. Our findings provide novel insights into how sex influences hippocampal neurogenesis and cognition in aging, and to highlight the relevance of considering biological sex in the design of therapeutic strategies for age-related cognitive decline.

## METHODOLOGY

### Compound and Treatment

The isolation and purification of 12-deoxyphorbol 13-isobutyrate—also known as DPB (Ezzanad et al., 2021) or ER272 (Dominguez-Garcia et al., 2021) (CAS: 25090-74-8) were carried out in our laboratory according to the procedures described in the supplementary methods of Geribaldi-Doldan et al. (Geribaldi-Doldan et al., 2015).

### Animal Subjects

Male and female SAMR1 and SAMP8 mice, aged four months, were housed in a temperature-controlled environment (21–23°C) with a 12-hour light/dark cycle and had ad libitum access to standard chow (AO4 maintenance diet, SAFE, Épinay-sur-Orge, France) and water. All procedures involving animals adhered to the European Union Directive (2010/63/EU) and the Spanish legal frameworks (65/2012 and RD53/2013) regulating the use of animals in research. Experiments were reported in accordance with the ARRIVE guidelines (Kilkenny et al., 2010)

The estrous cycle of female mice was not monitored during the experimental period. Given the two-month chronic treatment, transient hormonal variations associated with the cycle were considered unlikely to exert a consistent or directional influence on the measured outcomes. Moreover, regular monitoring of the estrous cycle would have introduced additional handling-related stress, potentially confounding behavioral and molecular analyses. This approach is consistent with previous studies using long-term interventions in female rodents in which the estrous stage was not a primary experimental variable (Prendergast, Onishi, & Zucker, 2014).

### Treatments and experimental groups

Four-month-old male and female SAMP8 mice were treated with either vehicle or ER272 using intranasal administration for eight consecutive weeks (see below for details). Age-matched SAMR1 mice of both sexes were included as controls as shown in previous studies (Takeda, 2009). Throughout the treatment period, all animals received intraperitoneal injections of 5-bromo-2’-deoxyuridine (BrdU) every other day. The SAMR1 strain was used as a reference group to distinguish between baseline values and those related to the pathological features of the SAMP8 aging model. ER272 was administered exclusively to SAMP8 mice in order to evaluate its potential to prevent the cognitive and neurogenic impairments characteristic of this model. Similar experimental approaches have been previously reported (Iwata et al., 2020; Shi et al., 2020).

### Intranasal delivery of ER272

The compound ER272 was administered intranasally with the animals in vertical position and their necks gently extended, following procedures described by Marks et al. (Marks, Tucker, Cavallin, Mast, & Fadool, 2009) . A total of 18 µL ER272 solution (1 µM in 0.9% saline) or saline was dispensed into the nostrils, alternating 3 µL per nostril using a micropipette. Mice remained in this position for approximately 10 seconds post-administration to promote full nasal absorption. Treatment assignment (vehicle or ER272) was randomized and coded, and all outcome measures were evaluated by investigators blinded to group allocation.

### Experimental strategy for testing the effect of ER272 in neurogenesis and cognitive performance of SAMP8 mice

A schematic of the experimental workflow is shown in Figure 1A. Male SAMR1 and SAMP8 mice, four months of age, received intranasal applications of ER272 or vehicle for a period of eight weeks. SAMR1 animals served as the reference group as previously established (Takeda, 2009). Throughout the treatment, animals were injected intraperitoneally with BrdU every other day. During week 6, mice underwent behavioral assessments including the Morris Water Maze (MWM), novel object discrimination (NOD), and open field test. Animals were euthanized at the end of the eighth week.

**Figure 1.**
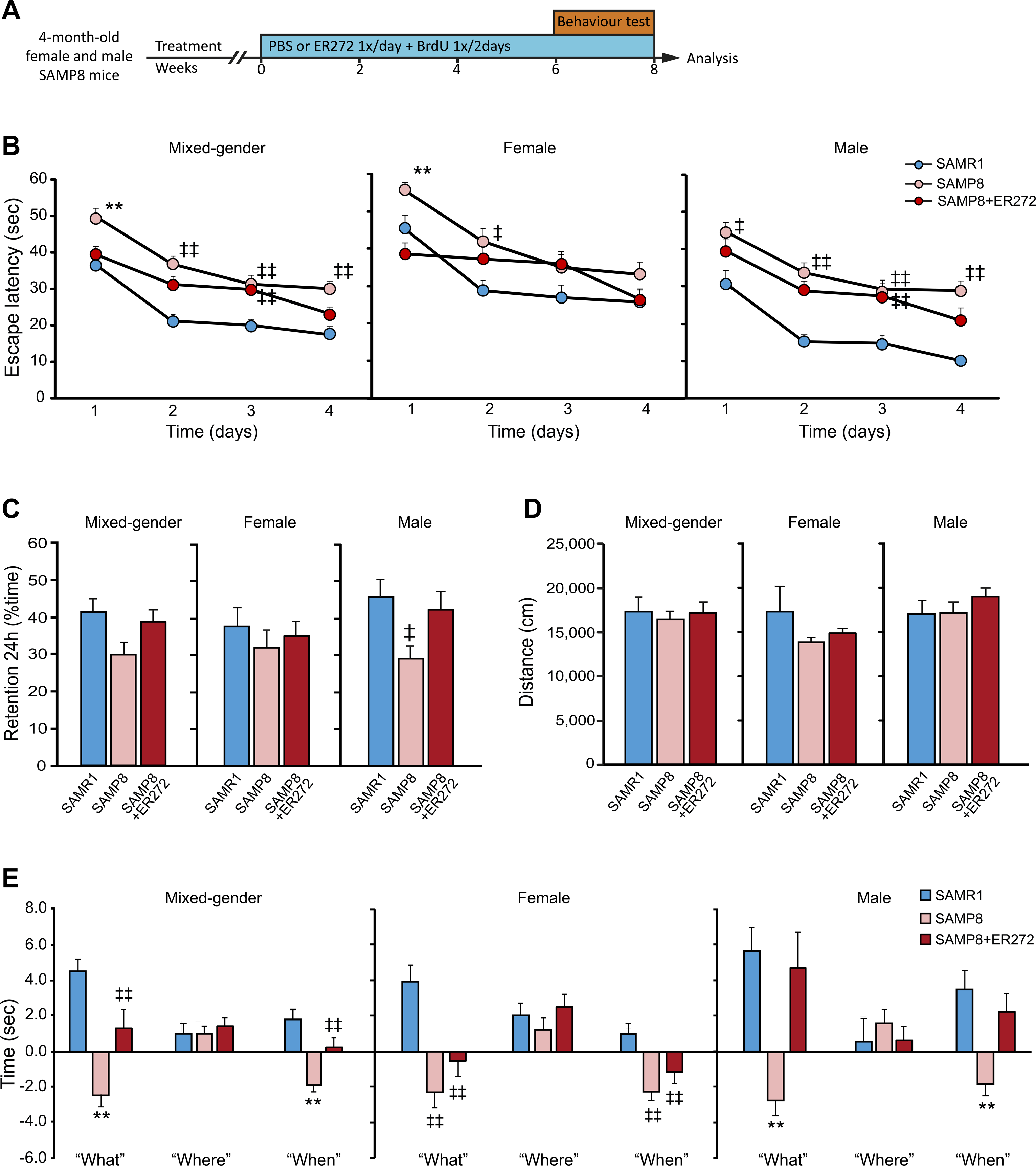
Treatment of SAMP8 mice with ER272 improves cognitive performance in male but not in female. **A) Experimental design.** Four-month-old male and female SAMP8 mice were treated with vehicle or ER272 (SAMP8+ER272) for 8 weeks until the age of 6 months. They were compared with SAMR1 mice of the same age treated with vehicle. Mice received BrdU during the 8-week period every two days. Behavior tests took place during the last two weeks. **B) Scape latency (sec) in the MWM test** (*Right*). A compromise was observed in SAMP8 male mice when they were evaluated in the MWM and a slight improvement was observed for SAMP8-ER mice. No significant group X day effect was detected [F(2,462)=0.358, p=0.905], however individual daily assessment revealed a better performance as training sessions progressed (day 1 [F(2,116)=4.64, ‡p=0.031vs. SAMR1], day 2 [F(2,116)=10.26, ‡‡p<0.01 vs. SAMR1], day 3 [F(2,113)=3.67, ‡‡p<0.001 vs. SAMR1], day 4 [F(2,116)=11.14, ‡‡p<0.001 vs. SAMR1]. Data are representative of 10 mice (SAMR1 n=10, SAMP8 n=10, SAMP8-ER n=10). (*Center*) A compromise was observed in SAMP8 female mice when they were evaluated in the MWM and a slight improvement was observed for SAMP8-ER mice. No significant group X day effect was detected [F(2,495)=2.02, p=0.061], however individual daily assessment revealed a better performance as training sessions progressed (day 1 [F(2,125)=8.75, **p<0.001 vs. rest of the groups], day 2 [F(2,126)=3.90, ‡p=0.023 vs. SAMR1], day 3 [F(2,115)=1.97, p=0.143], day 4 [F(2,129)=1.73, p=0.180]. Data are representative of 10-12 mice (SAMR1 n=10, SAMP8 n=11, SAMP8-ER n=12). **(Left)** A compromise was observed in SAMP8 mice when male and female cohorts were evaluated together in the MWM and a slight improvement was observed for SAMP8-ER mice. No significant group X day effect was detected [F(2,969)=1.898, p=0.078], however individual daily assessment revealed a better performance as training sessions progressed (day 1 [F(2,244)=9.21,**p<0.001 vs. rest of the groups], day 2 [F(2,245)=12.84, ‡‡p<0.001 vs. SAMR1], day 3 [F(2,231)=7.98, ‡‡p<0.001 vs. SAMR1], day 4 [F(3,249)=9.02, ‡‡p<0.001 vs. SAMR1]. Data are representative of 20-22 mice (SAMR1 n=20, SAMP8 n=21, SAMP8-ER n=22). **C) Retention test in the MWM.** In the 24 h retention phase SAMP8 male mice (right) spent shorter times in quadrant 2 where the platform used to be located. [F(2,25)=4.16, ‡p=0.028 vs. SAMR1]. No differences were detected among females (center) [F(2,29)=0.288, p=0.752]. No differences were observed in the retention phase when both cohorts were compared together (left) 24 h [F(2,58)=2.51, p=0.09] after completing the acquisition phase. **D) Distances travelled in the open field.** No differences were observed among groups when distances travelled in the open field were analyzed in males [F(3,27)=0.717, p=0.497], females [F(2,24)=0.89, p=0.420] or both cohorts together [F(2,54)=0.27, p=0.763]. Data are representative of 9-12 animals (males: SAMR1 n=10, SAMP8 n=10, SAMP8-ER n=10; females: SAMR1 n=8, SAMP8 n=12, SAMP8-ER n=7). Sex comparisons: motor activity (total distance travelled) in males and females in the groups under study (SAMR1 [F(1,16)=0.845, p=0.372], SAMR1-ER [F(1,18)=2.62, p=0.122], SAMP8 [F(1,15)=3.55, p=0.560], SAMP8-ER [F(1,20)=1.40, p=0.250]). Differences detected by one-way ANOVA followed by Tuckey b test. **E) NOD test.** (*Right*) Episodic memory was severely affected in male SAMP8 mice when “what” and “when” paradigms were assessed, while ER treatment counterbalanced this situation. “What” [F(2,84)=9.55, **p<0.001 vs. rest of the groups], “where” [F(2,84)=0.309, p=0.735], “when” [F(2,89)=8.69, **p<0.001 vs. rest of the groups]. (*Center*) Episodic memory was severely affected in female SAMP8 mice when “what” and “when” paradigms were assessed, while ER treatment improved this impairment, differences did not reach statistical significance. “What” [F(2,93)=15.60, ‡‡p<0.001 vs. SAMR1 ], “where” [F(2,96)=1.11, p=0.332], “when” [F(2,90)=8.81, ‡‡p<0.001 vs. SAMR1 ]. (*Left*) Episodic memory was severely affected in SAMP8 mice when “what” and “when” paradigms were assessed in both male and female cohorts together, while ER treatment counterbalanced this situation. “What” [F(2,180)=20.63, **p<0.001 vs. rest of the groups, ‡‡p<0.001 vs. SAMR1], “where” [F(2,183)=0.132, p=0.876], “when” [F(2,180)=15.33, **p<0.001 vs. rest of the groups, ‡‡p<0.001 vs. SAMR1]. Sex comparisons: in the NOD test SAMP8-ER females showed worse responses in the “what” and “when” paradigms than males (what: SAMR1 [F(1,58)=0.541, p=0.465], p=0.898], SAMP8 [F(1,58)=0.030, p=0.864], SAMP8-ER [F(1,61)=7.161, p=0.01]; where: SAMR1 [F(1,55)=1.265, p=0.266], SAMP8 [F(1,61)=0.100, p=0.753], SAMP8-ER [F(1,64)=3.776, p=0.056]; when: SAMR1 [F(1,58)=2.456, p=0.068], p=0.732], SAMP8 [F(1,61)=0.836, p=0.364], SAMP8-ER [F(1,58)=8.305, p=0.006]).

### Evaluation of locomotor activity and NOD

Behavioral testing began ten days before the end of the study. Locomotor activity was quantified by tracking the distance moved during a 30-minute interval prior to NOD testing. The habituation phase began with the exposure to two different objects (a red cylinder and a yellow trapezoid). One day after the habituation phase, we assessed integrated episodic memory for the paradigms “what,” “when,” and “where” (Ramos-Rodriguez et al., 2013). On day one, mice explored the habituation objects. On the following day, they encountered four copies of a new object (blue balls) arranged in a triangular pattern for 5 minutes. After 30 minutes, a second exposure included four novel items (red cones) in a square configuration. The final test phase, conducted 30 minutes later, involved two “recent” objects from the second exposure (in unchanged positions) and two “familiar” objects from the first exposure—one in the original location and one relocated. Memory components were quantified as: “what” (exploration time difference between familiar and recent objects), “where” (difference in time spent with displaced vs. non-displaced objects), and “when” (preference between familiar and recent items in original positions), following previously established experimental designs (Dere, Huston, & De Souza Silva, 2005; Infante-Garcia et al., 2018; Segado-Arenas et al., 2018).

### MWM

Forty-eight hours after completing the NOD task, spatial learning and memory performance were evaluated using the MWM test in all experimental groups, based on the procedure described by Ramos-Rodriguez et al. (Ramos-Rodriguez et al., 2013). In brief, the Morris water maze consisted of a circular tank measuring 95 cm in diameter, virtually divided into four quadrants. Visual cues in the form of geometric figures were placed around the pool to aid spatial orientation. The escape platform was positioned in quadrant 2 and submerged 2 cm below the water surface, with water maintained at 211±11 °C. The acquisition phase spanned four consecutive days, during which each mouse underwent four trials per day, spaced 10 minutes apart. Each trial had a maximum duration of 60 seconds, and mice were allowed to remain on the platform for 10 seconds upon reaching it. If the animal failed to locate the platform within the allotted time, it was gently placed on it for the same duration. Twenty-four hours after the last training session, a probe test was conducted: the platform was removed, and mice were allowed to swim freely for 60 seconds. Escape latency during the acquisition trials and the time spent in the target quadrant during the probe test were measured and analyzed using SMART software (Panlab, Spain).

### Tissue processing and immunohistochemistry

At the study’s end, brains were fixed via PFA perfusion and sectioned at 30 µm thickness using a cryostat. Immunostaining followed protocols described in earlier publications (Garcia-Bernal et al., 2018; Geribaldi-Doldan et al., 2015; Murillo-Carretero et al., 2017). Detailed antibody information is provided in Supplementary Tables 1 and 2.

### Quantification of hippocampal neurogenesis

BrdU, doublecortin (DCX), NeuN, glial fibrillary acidic protein (GFAP), SRY-box transcription factor 2 (SOX2), and S100 calcium-binding protein B (S100β)-positive cells in the DG were quantified as previously outlined (Rabaneda et al., 2008; Rabaneda et al., 2016). After perfusion, brains were anonymized with coded labels. Cell counts were performed blindly on every fifth coronal section encompassing the hippocampus (14–16 sections per animal), using a Zeiss LSM 900 Airyscan 2 confocal microscope. Z-stacks were acquired every 1.5 μm with a 20× objective. Cell density values were normalized to the DG volume and averaged per animal, as described by Dominguez-Garcia et al. (Dominguez-Garcia et al., 2021).

### Morphometric analysis of DCX-positive neurons

Three-dimensional reconstructions of DCX-labeled neurons were obtained from confocal image stacks using Neurolucida 360 software (MBF Bioscience, VT, USA). Morphological parameters, including total dendritic length, surface area, segment number, terminal branches, and Sholl intersections, were quantified using Neurolucida Explorer (Carrascal, Nieto-Gonzalez, Torres, & Nunez-Abades, 2009; Nunez-Abades, He, Barrionuevo, & Cameron, 1994).

### Statistical approach

All statistical analyses followed pharmacological research guidelines for rigorous experimental design (Curtis & Abernethy, 2015). Analyses were conducted in IBM SPSS Statistics v22. Data normality was tested using the Shapiro-Wilk test, followed by Brown-Forsythe’s test for homogeneity of variances. One-way ANOVA with Tukey’s post hoc correction was used for multi-group comparisons. For the MWM acquisition phase, two-way ANOVA (group × day) was applied. Significance was set at p < 0.05. Sample sizes ranged from n = 6–10 (in vivo), n = 5–9 (in vitro), and n = 10–12 (behavioral), consistent with prior work (Carrasco et al., 2014; Dominguez-Garcia et al., 2020; Geribaldi-Doldan et al., 2018; Rabaneda et al., 2016).

## RESULTS

### Sex-specific efficacy of diterpenoid treatment: cognitive recovery in male but not female SAMP8 mice

To test whether adult hippocampal neurogenesis and cognitive performance differ between male and female SAMP8 mice and to determine their responsiveness to ER272 treatment, we used the experimental approach represented in figure 1A. Four-month-old male and female SAMP8 mice were treated with either vehicle or ER272 during eight consecutive weeks. Male and female SAMR1 mice were used as the control group according to previous reports (Takeda, 2009). Throughout the treatment period, mice received intraperitoneal injections of BrdU every other day. Two weeks before treatment completion, animals underwent behavioral testing (MWM, NOD, and open field). Finally, mice were euthanized at the end of the 8-week treatment period once behavioral test were completed.

Spatial memory was analyzed in six-month-old SAMP8 male and female mice after treatment with diterpene ER272, or vehicle. A compromise was observed in SAMP8 male and female mice when they were evaluated in the training phase of the MWM and a slight improvement was observed for SAMP8 male and female mice treated with ER272 (Fig.1 B). No significant group X day effect was detected. However individual daily assessment revealed a better performance of SAMP8 male and female treated mice as training sessions progressed comparable to that of SAMR1 mice. In the retention phase, SAMP8 male mice showed a statistically significant reduction of the time spent in the platform quadrant (Fig. 1 C). This reduction was not observed in female mice (Fig. 1C). Moreover, treatment with ER272 improved retention in male mice, whereas no differences were detected in female SAMP8 mice treated with ER272 compared to female SAMP8 treated with vehicle (Fig 1C). When we compared males and females from different groups in the acquisition phase of the MWM we did not detect a significant day X sex effect in any of the groups under study, although significant differences were observed for day and sex criteria (Suppl. Table 3). On the other hand, sex comparisons in the 24h or 72 h retention phases did not reveal significant differences in any of the groups under study. Similarly, no differences were detected when the number of entries into quadrant 2 were compared between sexes in all groups under study. As control, no differences were found in any of the groups when the distance travelled in the open field was analyzed (Fig. 1 D) and neither when male and female were compared.

Episodic memory was tested in the NOD test. Results showed that episodic memory was severely affected in male and female SAMP8 mice when “what” and “when” paradigms were analyzed (p<0.001). Treatment with ER272 rescued this situation only in male mice whereas no improvement was observed in female in response to the treatment (Fig. 1 E). When sexes were compared in the NOD test we observed that SAMP8-ER females had worse responses in the “what” and “when” paradigms than males.

### Sex-dependent differences in neurogenesis in the DG and their response to treatment in six-month-old SAMP8 mice

To investigate neurogenesis in the DG (Fig. 2 B) of SAMP8 mouse and assess the influence of sex and diterpene treatment, mice received BrdU injections every other day throughout the treatment period. As shown in Figure 2 A, C, both male and female SAMP8 mice exhibited a reduction in the number of BrdU-incorporating cells after two months of treatment, although this decrease did not reach statistical significance. Notably, treatment with ER272 led to a significant increase in BrdU^+^ cells in both sexes (Fig 2, A, C). This effect was particularly marked in male SAMP8 mice, where the number of BrdU^+^ cells exceeded that observed in SAMR1 controls (Fig. 2 A, C). Interestingly, the number of BrdU^+^ cells in ER272-treated males doubled that found in treated females (Fig. 2 A, C).

**Figure 2.**
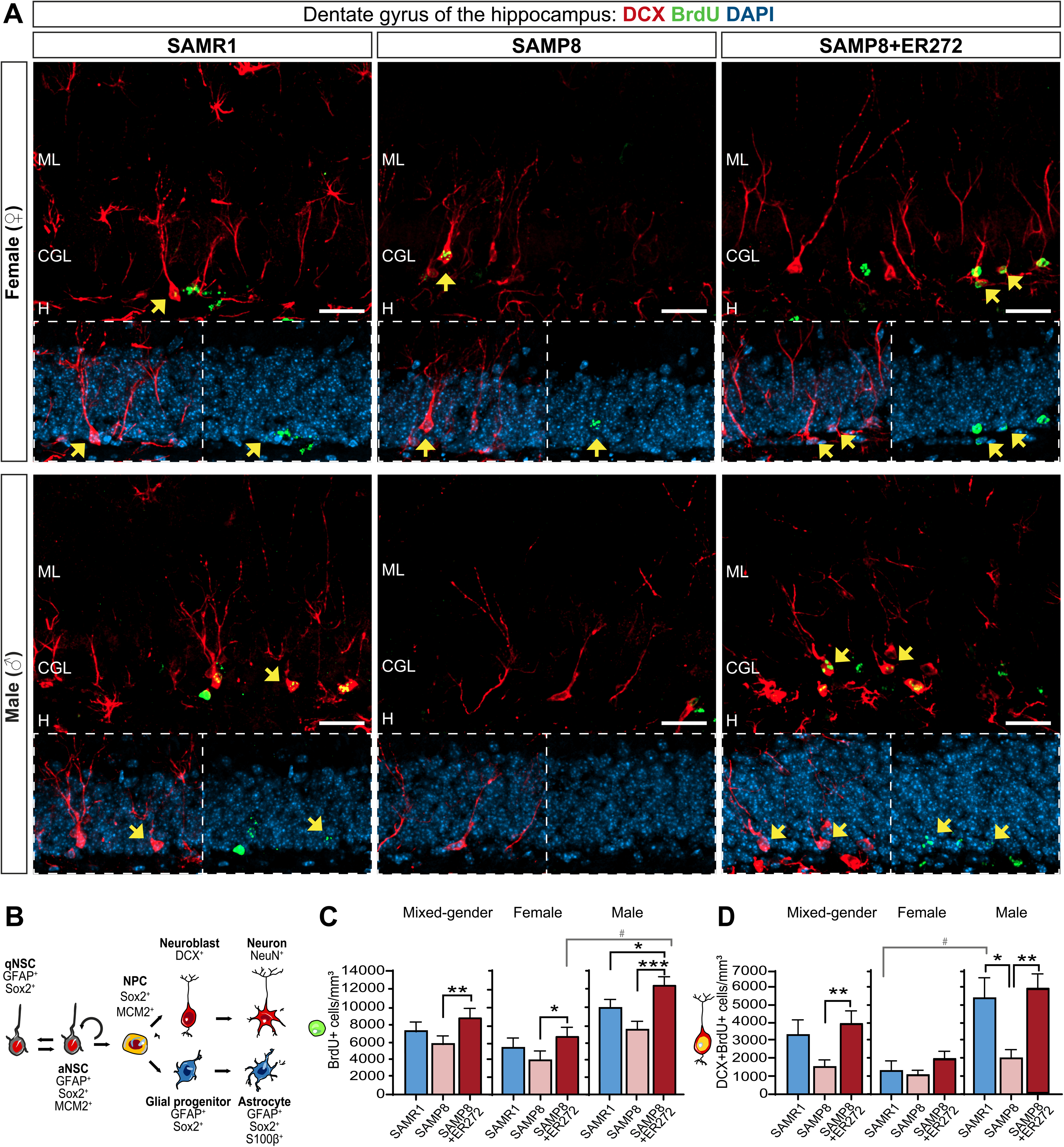
Two-month intranasal administration of ER272 to SAMP8 mice differentially affects DCX neuroblasts in male and female. **A)** Representative confocal microcopy images of the DG of the hippocampus of six-month-old SAMR1 (left) and SAMP8 male mice (center) treated with vehicle or SAMP8 mice treated with ER272 (SAMP8+ER272) right in female (upper panel) and male (lower panel) mice. Tissue was processed for the detection of the proliferation marker BrdU and DCX. DAPI staining is shown in blue. Only merged channels are shown. Yellow arrows indicate DCX^+^BrdU^+^DAPI^+^ cells. **B)** Schematic drawing of the hierarchical adult DG neurogenesis indicating the markers used to detect the progeny of NSCin the DG. **C)** Graph shows the total number of BrdU^+^ nuclei in the DG of the hippocampus per mm^3^ in male (right) [F =10.58, *p=0.045 SAMR1 vs SAMP8+ER272] [F =10.58, ***p<0.001 SAMP8 vs P8+ER272], female (center) [F_(3,51)_=3.94, *p=0.037 SAMP8 vs SAMP8+ER272] and both cohorts together (left) [F_(2,60)_=6.82, **p=0.001 SAMP8 vs SAMP8+ER272]. Sex differences in BRDU⁺ cells are also shown for SAMP8+ER272 group [F =11.2, # p=0.005] **D)** Graph shows the total number of BrdU^+^DCX^+^ cells in the DG of the hippocampus per mm^3^ in male (right panel) [F =6.898, *p=0.034 SAMR1 vs P8] [F =6.898, **p<0.003 P8 vs SAMP8+ER272], female (**center panel**) and both sexes together (left panel) [F_(2,57)_=4.44, **p=0.01 SAMP8 vs SAMP8+ER272]. Sex differences in DCX⁺BRDU⁺ cells are also shown for SAMR1 group [# p=0.02]. Data are the means⍰±⍰S.E.M of six animals, n⍰=⍰6. Differences detected by one-way ANOVA followed by Tukey b test. Scale bar represents 25 μm.

Analysis of immature neurons (DCX^+^ cells) generated over the course of the treatment (BrdU^+^/DCX^+^), indicated that the number of neuroblast during the treatment period (DCX^+^/BrdU^+^) increased in male SAMR1 by 5-fold compared to SAMR1 females. Interestingly, male SAMP8 mice showed a significant reduction in DCX^+^/BrdU^+^ cells, which was fully prevented by ER272 treatment (Fig. 2, A, D). In contrast, female SAMP8 mice did not differ from female SAMR1 controls in the number of newly generated DCX^+^/BrdU^+^ cells. This result indicated that control male generated a larger number of new neurons from months 4 to 6 compared to female. This is not observed is not in SAMP8 mice. On the contrary, the number of DCX^+^/BrdU^+^ cells in SAMP8 male mice is comparable to that of control and SAMP8 female. Remarkably, the treatment prevents this reduction.

We next analyzed the number of newly generated mature neurons (NeuN^+^/BrdU^+^ cells). Control SAMR1 male mice showed a higher number of mature neurons generated over the course of the treatment compared to female SAMR1 mice however, these differences did not reach statistical significance. No statistically significant differences were found between SAMP8 and SAMR1 mice in either sex (Fig. 3 A, B). However, ER272 treatment significantly increased the number of newly generated mature neurons in male SAMP8 mice, a tendency that was also observed in females, but it was not statistically significant.

**Figure 3.**
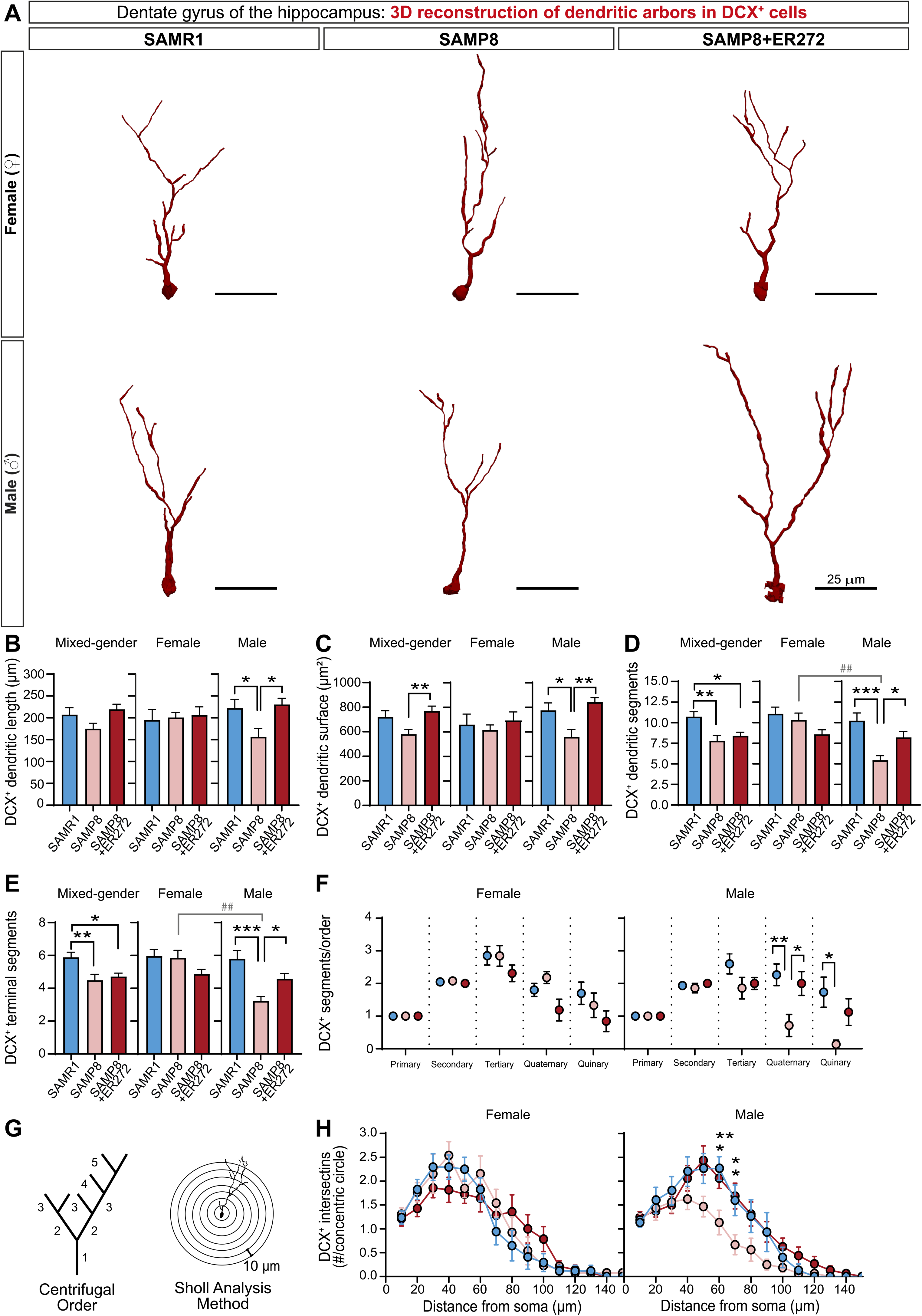
Sex dependent effect of intranasal administration of ER272 to SAMP8 mice on NeuN cells. **A)** Representative confocal microcopy images of the DG of the hippocampus of six-month-old SAMR1 and SAMP8 male mice treated with vehicle or SAMP8 mice treated with ER272 during eight weeks. Female (upper panel) and male (lower panel). Slices were processed for the immunohistochemical detection of the proliferation marker BrdU and NeuN. DAPI staining is shown in blue. Only merged channels are shown. Yellow arrows indicate NeuN^+^BrdU^+^DAPI^+^ cells. **B)** Graph shows the number of NeuN^+^BrdU^+^ nuclei in the DG of the hippocampus per mm^3^ in male (right) [F_(2,8)_=11.04, **p=0.04 P8 vs SAMP8+ER272], female (center) and both sexes together.

### Sex-specific alterations of the complexity of newly generated neurons within the neurogenic niche and treatment response in 6-month-old SAMP8 mice

The morphological properties of immature were then analyzed across all experimental groups. No differences were observed in total dendritic length between male and female SAMR1 control mice (∼ 200 µm) (Fig. 4 A, B). Male SAMP8 mice, however, showed a reduced dendritic length compared to SAMR1 controls, an effect that was prevented by ER272 treatment (Fig. 4 A, B). In contrast, SAMP8 females showed neither a reduction in dendritic length and nor effect of the treatment (Fig. 4 A, B). Dendritic surface area was also reduced in male SAMP8 mice relative to SAMR1 males, whereas no changes were detected in females. ER272 administration prevented this reduction in males (Fig. 4 A, C). Similarly, the number of dendritic segments and the number of terminal segments was reduced in male SAMP8 mice compared to SAMR1 but not in females (Fig. 4 A, D, E). Treatment with ER272 prevented this reduction in males whereas no effect of the treatment was observed in females (Fig. 4 A, D, E). In addition, segment order analysis showed a reduction in the number of quaternary and quinary segments in males that was prevented by ER272 in SAMP8 male mice. No statistically significant differences or treatment effect were detected in females (Fig. 4F,G). Finally, Sholl analysis showed a lower number of intersections at 60-70 μm from the soma in male SAMP8 mice compared to SAMR1 controls (Fig. 4 A, G-H), an effect that was reverted by ER272 treatment. Interestingly, no differences were observed in females (Fig. 4 A, G-H).

**Figure 4.**
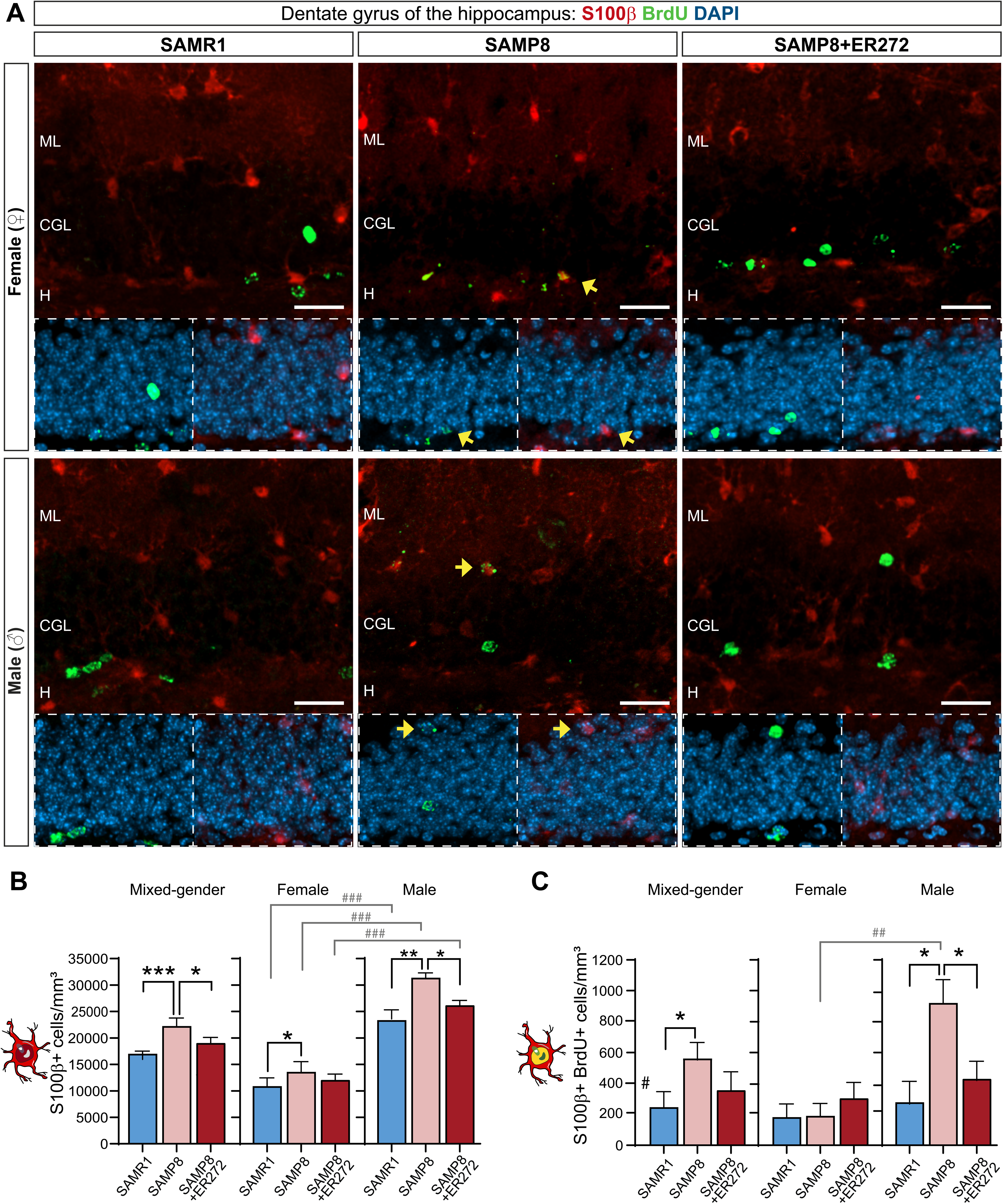
Sex dependent alteration of the morphology of DCX^+^ cells in SAMP8 and differential response to the treatment. **A)** Examples of the reconstructed DCX^+^ cells used in the analysis of SAMR1 and SAMP8 treated with vehicle and SAMP8 treated with ER272. Female (upper panel) and male (lower panel). **B)** Total dendritic length of females, males [F_(2,43)_=4.94, *p=0.040 SAMR1 vs SAMP8; *p=0.016 SAMP8 vs SAMP8+ER272] and both sexes. **C)** Total dendritic surface of females, males [F_(2,43)_⍰=⍰7.157, *p⍰=⍰0.024 SAMR1 vs. SAMP8; **p⍰=⍰0.002 SAMP8 vs. SAMP8+ER272]] and both sexes [**p=0.003 SAMP8 vs SAMP8+ER272]. **D)** Number of dendritic segments of females, males [F_(2,40)_=9.51, ***p<0.001 SAMR1 vs SAMP8; *p=0.038 SAMP8 vs SAMP8+ER272] and both sexes [**p=0.003 SAMP8 vs SAMP8+ER272; *p=0.03 SAMR1 vs SAMP8+ER272]. Sex differences in dendritic segments are also shown for the SAMP8 group ( [ ## p<0.002]**E)** Number of terminal segment of females, males [F_(2,41)_=10.5, ***p<0.001 SAMR1 vs SAMP8; *p=0.045 SAMP8 vs SAMP8+ER272] and both sexes [**p=0.005 SAMR1 vs SAMP8; *p=0.03 SAMR1 vs SAMP8+ER272]. Sex differences in terminal segments are also shown for the SAMP8 group [ ## p<0.002] **F-G)** Number of segments per order of females, males (Quaternary order [F_(2,42)_⍰=⍰5.50, **p⍰<⍰0.009 SAMR1 vs. SAMP8] [F_(2,42)_⍰=⍰5.50, *p⍰=⍰0.032 SAMP8 vs. SAMP8+ER272]; Quinary order [F_(2,43)_⍰=⍰4.76, *p⍰<⍰0.011 SAMR1 vs. SAMP8]). **G-H)** Sholl dependent analysis of females and males (60⍰μm from soma [F_(2,43)_⍰=⍰6.292, **p⍰<⍰0.005 SAMR1 vs. SAMP8; *p⍰<⍰0.022 SAMP8 vs. SAMP8+ER272]; 70⍰μm from soma [F_(2,42)_⍰=⍰4.917, *p⍰<⍰0.026 SAMR1 vs. SAMP8; *p⍰<⍰0.025 SAMP8 vs. SAMP8+ER272]).

### Differential impact of sex on niche astrocytes and treatment efficacy in middle-aged SAMP8 mice

Disproportionate astrogliogenesis in the DG niche is a characteristic of brain aging (Encinas et al., 2011). To assess whether sex had an impact on these alterations, we analyzed astrogliogenesis in senescent mice and evaluated the effect of the ER272 diterpene treatment on the generation of new astrocytes. As expected, a larger number of niche astrocytes (GFAP^+^S100β^+^ cells) were found in male and female SAMP8 mice compared to male SAMR1. In both sexes the treatment prevented the enhanced astrogliogenesis (Fig. 5 A, B). Interestingly, the number of niche astrocytes in female mice of all groups was reduced by around 2-fold when compared to males (Fig. 5 A, B). The number of astrocytes generated during the two-month period (GFAP^+^/S100β^+^/BrdU^+^cells) was only found elevated in male SAMP8 mice by almost 4-fold compared to male SAMR1. This effect was prevented by the two-month ER272 treatment. On the contrary no differences were found in female SAMP8 mice compared to SAMR1 (Fig. 5 A, C). Interestingly, the number of GFAP^+^/S100β^+^/BrdU^+^cells in male SAMP8 mice was around 5-fold higher than in SAMP8 female. Indicating that astrogliosis is highly elevated in SAMP8 male mice compared to female.

**Figure 5.**
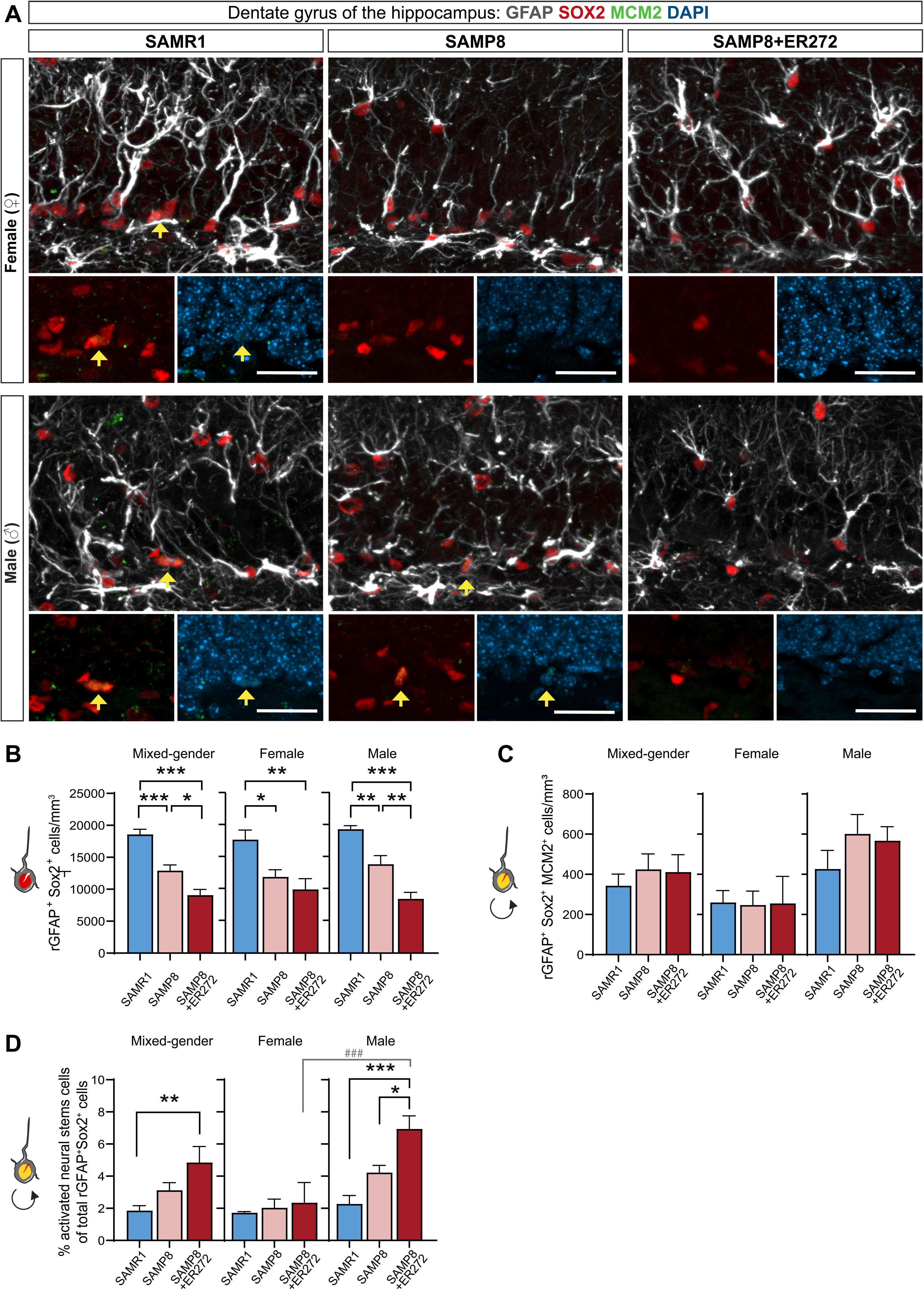
Sex differences in newly generated astrocytes in the DG of SAMP8 mice. **A)** Representative confocal microcopy images of the DG of the hippocampus of six-month-old SAMR1 and SAMP8 male mice treated with vehicle or SAMP8 mice treated with ER272 (SAMP8+ER272) for eight weeks as indicated in figure 1 A. Slices were processed for the immunohistochemical detection of the proliferation marker BrdU (green), the Glial the marker of mature astrocytes S100**β** (red) and DAPI. Yellow arrows indicate S100β^+^BrdU^+^ cells. **B)** Graph shows the number of S100β^+^ cells in the DG per mm^3^ in male [F =8.824, **p=0.001 SAMR1 vs SAMP8] [F =8.824, *p<0.041 SAMP8 vs P8+ER272], female [F_(2,19)_=10.58, *p=0.039 SAMR1 vs SAMP8] and both sexes together [F_(2,45)_=8.877, ***p<0.001 SAMR1 vs SAMP8] [F =8.877, *p<0.025 SAMP8 vs SAMP8+ER272] # Sex differences in the number of S100β^+^ cells in the DG per mm^3^ are also shown for all groups [F =29.9 ### p<0.001 SAMP8; ### p<0.001 SAMR1; ### p<0.001 SAMP8+ER272] **C)** Graph shows the number of S100β^+^BrdU^+^ cells in the DG of the hippocampus per mm^3^in male [F_(2,13)_=5.5605, *p=0.020 SAMR1 vs SAMP8] [F_(2,13)_=5.5605, *p<0.049 SAMP8 vs SAMP8+ER272], female and in both sexes together [*p=0.032 SAMR1 vs SAMP8].

### Sex-specific activation of niche NSCs in aged SAMP8 mice

The pool of NSCs and their activation was then analyzed. To this end, we studied the number of cells with radial glial like morphology expressing glial marker protein glial fibrillary acidic protein (GFAP) and transcription factor SOX2 (rGFAP^+^SOX2^+^). 6-month-old male and female SAMP8 mice showed a reduction in the number of rGFAP^+^SOX2^+^ cells. The treatment induced a further reduction in the number of these cells only in male, but not in female (Fig. 6 A, B). The analysis of the results including mice of both sexes render results similar to those found in males. Interestingly, male SAMP8 mice showed a higher number of activated NSC (rGFAP^+^SOX2^+^MCM2^+^) compared to females (Fig. 6 A, C). However, no alterations in the number of (rGFAP^+^SOX2^+^MCM2^+^) were found in SAMP8 mice compared to SAMR1, in either males or females. Furthermore, no statistically significant effect of the treatment on rGFAP^+^SOX2^+^MCM2^+^ cells was observed (Fig. 6 A, C) in any of the groups. Interestingly, the percentage of activated NSC relative to the total NSC population was higher in male SAMP8 mice than in male SAMR1. This percentage was incremented in male SAMP8 mice as a consequence of the treatment (Fig. 6 A, D). In contrast, no changes in the percentage of activated NSC was observed in female mice (Fig. 6 A, D). The analysis using animals of both sexes showed results similar to those found in male mice (Fig. 6), emphasizing the importance of analyzing sexes separately in this type of studies.

**Figure 6.**
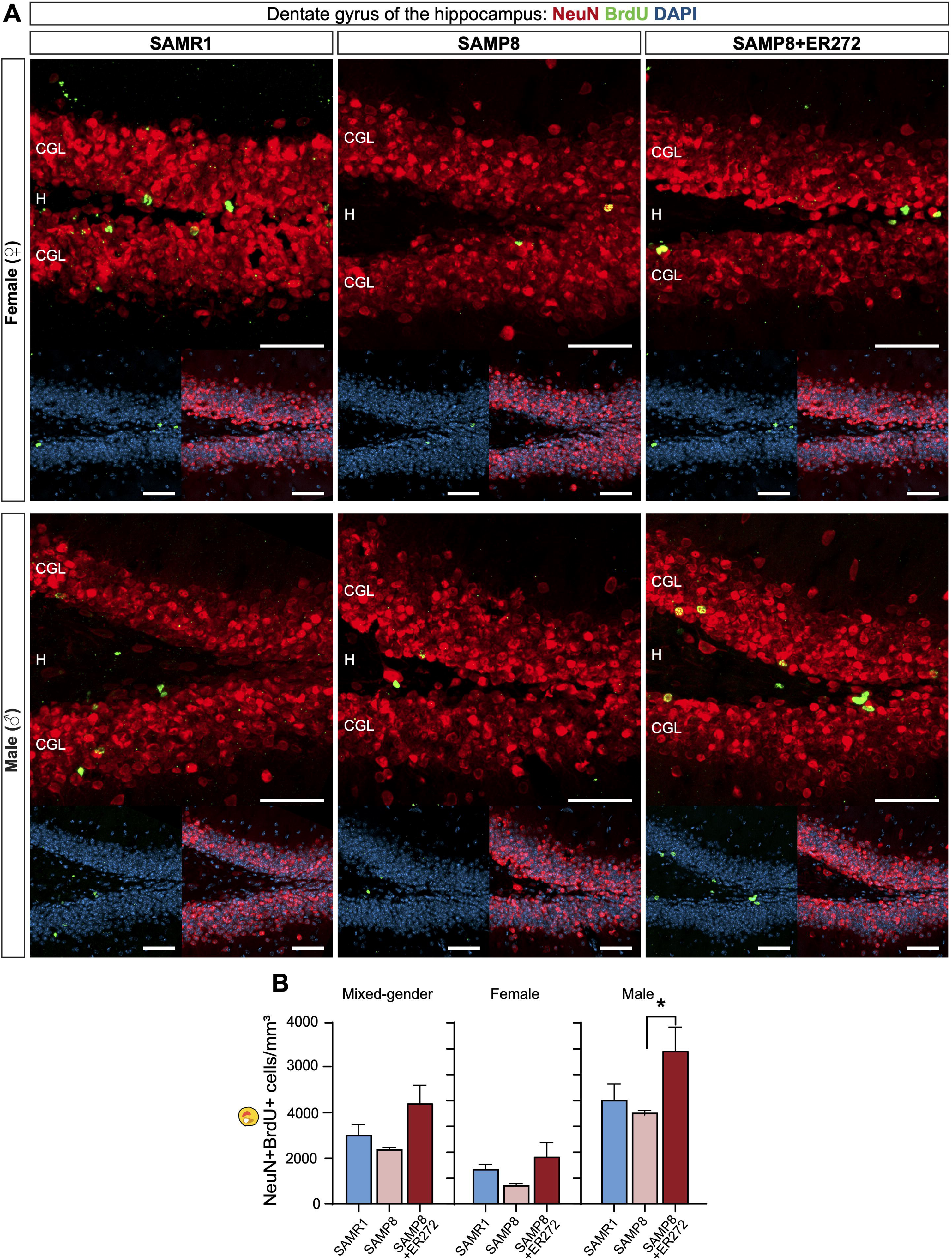
Sex dependent activation of NSC in the DG of SAMP8 mice and response to treatment. **A)** Representative confocal microcopy images of the DG of the hippocampus of six-month-old SAMR1 and SAMP8 male mice treated with vehicle or SAMP8 mice treated with ER272 during eight weeks as indicated in figure. 1 A. Tissue processed for the detection of the Glial Fibrillary Acidic Protein GFAP (white), SOX2 (red) and the cell cycle marker MCM2 (green). Yellow arrows indicate rGFAP^+^MCM2^+^SOX2^+^ cells. **B)** Graph shows the total number of rGFAP^+^SOX2^+^ with a radial glial-like cell (r) morphology in the DG per mm^3^ in female [F_(2,13)_=7.979, *p=0.023 SAMR1 vs SAMP8; **p<0.008 SAMR1 vs SAMP8+ER272], male [F_(2,15)_=27.92, **p=0.005 SAMR1 vs SAMP8; ***p<0.001 SAMR1 vs SAMP8+ER272; **p<0.005 SAMP8 vs SAMP8+ER272] and in both sexes together [F_(2,31)_=29.90, ***p<0.001 SAMR1 vs SAMP8; ***p<0.001 SAMR1 vs SAMP8+ER272; *p<0.012 SAMP8 vs SAMP8+ER272] **C)** Graph shows the number of rGFAP^+^SOX2^+^MCM2^−^ with a radial glial-like cell (r) morphology in the DG per mm^3^ in female, male and in both sexes. **D)** Percentage of activated NSC (rGFAP^+^SOX2^+^MCM2^+^) among the total number of NSC population (rGFAP^+^SOX2^+^) in the DG of females and males [F =14.3, ***p<0.001 SAMR1 vs SAMP8+ER272; *p<0.019 SAMP8 vs SAMP8+ER272] and in both sexes [F_(2,32)_=5.470, **p<0.007 SAMR1 vs SAMP8+ER272]. Sex differences in the percentage of activated NSC are also shown for the SAMP8+ER272 group [F_(5,28)_=8.73, ### p<0.001]

### Analysis of combined cohorts show a clear bias towards the male phenotype

Across the different parameters analyzed in this study (newly generated neurons, mature neurons, astroglial cells, NSCs, and cognitive performance), pooling data from both sexes resulted in values that were largely similar to those found in male mice. With the exception of the total number of NSCs, mixed-sex analyses failed to capture the female-specific patterns observed when sexes were analyzed separately.

## DISCUSSION

Cognitive performance tends to deteriorate with advancing age, and aging itself constitutes a major independent risk factor for neurodegenerative disorders affecting both sexes (Anstey et al., 2021). Certain memory domains, particularly episodic memory, appear to be especially vulnerable in older individuals (Gray & Barnes, 2015; Nyberg, Lovden, Riklund, Lindenberger, & Backman, 2012). As a result, identifying effective strategies to prevent or mitigate age-related cognitive decline has become a priority. Among the potential therapeutic approaches, potentiating adult hippocampal neurogenesis is a promising strategy to reduce cognitive deterioration in aging populations. However, to accelerate therapeutic discovery, it is crucial to examine sex differences. Experimental studies show that spatial training paradigms such as the MWM or pattern separation tasks selectively enhance DG neurogenesis of male but not female rats (Chow et al., 2013; S. Yagi & Galea, 2019). These sex-dependent effects on neurogenesis align with a superior behavioral outcome generally observed in males, raising the possibility that neurogenic responses may underlie at least part of the differences in cognitive performance found between sexes. We have previously identified molecules that enhance neurogenesis in the adult brain and proposed their potential as pharmacological interventions for cognitive deficits associated with impaired neurogenesis. Among these, the diterpene ER272, characterized by a 12-deoxyphorbol structure (Geribaldi-Doldan et al., 2015; Murillo-Carretero et al., 2017), was administered to male SAMP8 mice prior to the onset of cognitive decline, effectively preventing cognitive impairment. Extending these findings, our current results indicate that the beneficial effects observed in male SAMP8 mice do not necessarily generalize to both sexes. We show here that adult hippocampal neurogenesis differs between aged male and female SAMP8 mice, and that each sex exhibit distinctive responses to the treatment.

### Prevention of Cognitive Decline by Diterpenoid ER272 is Male-Specific

The behavioral analysis, using the MWM and NOD test, of 6-month-old SAMP8 mice treated for 2 months with diterpene ER272 revealed that although both male and female show a reduction in the capacity to perform these tasks at 6 months of age, only in male was this decline prevented by the treatment. Our results were consistent with previous findings on this model in which SAMP8 mice show cognitive decline by 6 months (Gomez-Oliva, Martinez-Ortega, et al., 2023; Pačesová et al., 2022). This suggested the existence of underlying sex-specific mechanisms leading to neurodegeneration or neuroprotection. Although several reports have shown a clear male advantage in spatial memory using different tests (Hausmann, Slabbekoorn, Van Goozen, Cohen-Kettenis, & Güntürkün, 2000; LaBuda, Mellgren, & Hale, 2002; Rodríguez, Chamizo, & Mackintosh, 2011), our results show no sex differences in performance in the NOD and MWM between control and SAMP8 mice, indicating no inherent advantage for either sex in this model at six months of age. However, we have observed a clear improvement in treated male suggesting a greater capacity of the male brain to respond to the treatment that correlates with neurogenesis during the treatment period.

### Sex-dependent differences in DG neurogenesis correlate with differences in cognitive performance

Sex differences have been reported in pattern separation, pattern completion, spatial learning, and their links to adult neurogenesis, highlighting the need to include both sexes when studying hippocampal cellular and structural mechanisms (S. Yagi & Galea, 2019). Here, we have evaluated separately DG neurogenesis in 6-month-old male and female SAMP8 mice. In addition, we have studied whether male and female respond differently to a two-month treatment with diterpene ER272.

We observe that the number of immature neurons generated from months 4-6 (BrdU^+^/DCX^+^) is higher in control SAMR1 male than in control SAMR1 female mice, suggesting the existence of a neurogenic response in males that is not present in female. However, SAMP8 male mice show a dramatic reduction in newly generated neurons, compared to male SAMR1, not found in SAMP8 female mice, which is prevented by the treatment. These results indicated that in males, neurogenesis is more active than in females from months 5 and 6 of life but this is reduced by aging. A reduction on the neurogenic response due to aging and neurodegeneration has been previously reported not only in murine models like the SAMP8 mice (Diaz-Moreno et al., 2018; Diaz-Moreno et al., 2013) or AD genetic models (Choi et al., 2018; Zaletel et al., 2018) but also in human (Marquez-Valadez, Rabano, & Llorens-Martin, 2022; Terreros-Roncal et al., 2021). In female neurogenesis was only comparable to that found in SAMP8 male mice suggesting that neurogenesis is reduced in control female mice as much as in aged SAMP8 female mice.

Remarkably, a neurogenic response was observed in control and in SAMP8 treated mice and they correlate with the results obtained in the cognitive tests in which, cognitive impairment was only prevented by the treatment in male and not in female. The analysis of neuronal maturation into NeuN^+^ neurons revealed that in male mice a higher number of newly generated neurons was found compared to female in all groups. Although these differences were not statistically significant, the tendency was clear. Also, in both male and female aged SAMP8 mice, a reduction in the number of mature neurons generated during months 5 and 6 was observed that was not statistically significant. Nevertheless, compound ER272 seems to promote maturation of newly generated cells in male mice and not in female. Previous works using male mice show an induction of proliferation in the young SAMP8 mice but a reduction in neurogenesis similar to that found in here in SAMP8 male mice at later stages (Diaz-Moreno et al., 2018; Diaz-Moreno et al., 2013).

### Sex-dependent differences in the maturation of new neurons correlate with cognitive performance

We found no statistically significant differences in the maturation of newly generated neurons between control SAMR1 male and female mice or between male and female SAMP8 mice. However, we observed that males were able to generate a higher number of neurons in response to the treatment compared to females.

It was interesting to notice several differences in the complexity of newly generated neurons. We observed no differences in the complexity of newly generated neurons between male and female control SAMR1 mice at the age of 6 months. However, the analysis of dendritic length, surface, segments and terminal segments in male SAMP8 mice revealed clear differences compared to male SAMR1 controls. These differences were not observed in female. Previous works using male SAMP8 mice show dysfunction and aberrant maturation of immature neurons in SAMP8 male mice that may correlate with our present findings (Diaz-Moreno et al., 2018; Diaz-Moreno et al., 2013). A previous report has studied the maturation of newly generated cells in the hippocampus of two month-old rats showing a slower differentiation rate in female than in male (S. Yagi et al., 2020). We do not observe this pattern in 6-month-old SAMR1 mice. However, we detect a clear delay in maturation in male SAMP8 mice compared to male SAMR1 mice that is prevented by the treatment. This difference in maturation may explain why SAMP8 male mice show a 4-fold reduction in the number of newly generated immature neurons compared to SAMR1 whereas the number of mature neurons remains similar. Previous reports have shown that newborn neurons that do not adequately complete maturation and synaptic integration programs are promptly eliminated in rodents (Kempermann, Jessberger, Steiner, & Kronenberg, 2004; Sierra et al., 2010) and that SAMP8 male mice have shown an increase in apoptosis correlating with the presence of aberrant neurogenesis (Moreno-Jiménez et al., 2019). It is possible that the reduced maturation capacity found in male SAMP8 DCX^+^ cells might underlie the marked decline in DCX^+^/BrdU^+^ cells observed in male SAMP8 mice compared to SAMR1. This phenomenon has been observed in human subjects (Moreno-Jiménez et al., 2019). This sharp decline is not observed in the number of NeuN^+^/BrdU^+^ cells suggesting that those that have reached maturation survived in SAMP8 male mice. The delay in maturation is not present in female SAMP8 mice. This indicates that the fewer newly generated DCX^+^/BrdU^+^ cells found in aged SAMP8 female become fully differentiated neurons whereas in SAMP8 mice these new neurons are aberrant, not fully mature and susceptible to undergo apoptosis.

The capacity of the treatment to promote neurogenesis in SAMP8 male mice correlates with the observations found in behavioral tests in which a greater capacity of male to respond to the treatment than female is found and emphasize the importance of including both sexes when testing hippocampal neurogenesis and spatial memory during aging.

### Age-dependent astrocytosis in the DG can be prevented in male but not in female with diterpene treatment

Previous reports have shown that hippocampal aging is characterized by the differentiation of NSCs into astroglial cells leading to the loss of NSC in the DG of aged mice due to their preferential differentiation into mature astrocytes (Diaz-Moreno et al., 2018; Encinas et al., 2011). In our study, we observe a higher number of astrocytes generated during the two-month treatment period in six-month-old SAMP8 male mice compared to SAMR1 controls. The generation of a higher number of astrocytes from month 4 to moth 6 was not observed in female. Nevertheless, the treatment prevented this reduction in male. It was also noticeable the increase in the number of total astrocytes in both sexes, although female showed a smaller number of astrocytes compared to male. These results are consistent with the reduced number of DCX^+^ cells generated during the treatment period in male compared to female. In line, suggesting that activated NSC in SAMP8 male mice are giving rise to astrocytes rather than neuroblasts. In addition, these results are consistent with the higher number of newly generated astrocytes found in male SAMP8 mice. Consistent with previous reports (Moreno-Jiménez et al., 2019), these findings suggest astrogliogenesis from NSC is favored in male SAMP8 mice, and that the treatment can prevent this condition specifically in male.

### Sex-dependent activation of NSCs

Evidence show that the proportion of NSC decreases with age in the mouse brain. Concomitanly, a reduced the ability of NSC to undergo activation is found that allows them to remain quiescent for prolonged periods of time (Diaz-Moreno et al., 2018; Harris et al., 2021; Ibrayeva et al., 2021; Martin-Suarez, Valero, Muro-Garcia, & Encinas, 2019). This extended quiescence avoids exhaustion of the NSC pool. We observed a smaller number of NSC in the DG of SAMP8 male and female mice consistent with previous reports (Diaz-Moreno et al., 2018) as demonstrated by the lower number of radial glial like cells that express SOX2 and GFAP. This number was even lower in treated SAMP8 male mice, but not in females. Interestingly, the percentage of activated NSC increases in SAMP8 male mice compared to SAMR1 and is higher in SAMP8 treated male mice than in non-treated SAMP8 male mice. This effect was not observed in females. Previous reports have found two different types of NSC in the aged mouse brain. NSC higher activation rate and greater contribution to neurogenesis and NSC with a much lower probability of division (Harris et al., 2021; Ibrayeva et al., 2021; Martin-Suarez et al., 2019). The different response in NSC activation found in SAMP8 male vs female might be explained by the presence in male of a higher number of resting, non-dormant cells (Harris et al., 2021; Martin-Suarez et al., 2019) that account for the higher mitotic activity found in male mice in response to the treatment and that contribute to a greater plasticity in the aged hippocampus. It is possible that prolonged treatment with this compound in male could eventually lead to depletion of the NSC pool.

### Sex Bias in Mixed Cohorts: Male-Driven Outcomes in Aging Studies

In most of our studies using mixed cohorts of SAMP8 mice, the resulting phenotype closely resembled that of males, rather than representing an intermediate or averaged response. This male bias in mixed analyses has been previously described in multiple preclinical studies, where male data often dominate due to greater variability or a stronger expression of age-related phenotypes. In agreement with previous reports, these findings reinforce the importance of considering sex as a biological variable in experimental design and data interpretation, in order to uncover mechanisms that may be unique to the female brain (Beery & Zucker, 2011; Shansky & Woolley, 2016).

## Conclusion

In conclusion, our results show that aged male and female SAMP8 mice perform similarly in behavioral tests assessing cognitive function; however, they exhibit distinct responses to diterpene treatment, with significant effects observed only in males. In male mice, the treatment enhanced neurogenesis and promoted the maturation of newly generated neurons, whereas this response was markedly reduced in females. Moreover, the NSC pool was more active in males and further stimulated by the treatment, suggesting a greater neurogenic response in the male dentate gyrus under these conditions.

These findings highlight the importance of including both sexes in studies evaluating therapeutic interventions, as results obtained from males or mixed cohorts may not accurately reflect female-specific responses.

## ACKNOWLEDGMENTS

We thank the “Servicio de experimentación y producción animal (SEPA) de la Universidad de Cádiz” as well as the “Servicios Centrales de apoyo a la investigación en BIOMEDICINA (SCIBM)” and “Servicios centrales de Ciencia y tecnología (SC-ICYT)” de la Universidad de Cádiz.

