## supplementary information for "Sex-Specific neurogenic and cognitive responses in a murine model of accelerated aging"

**Supplementary table 1. Primary antibodies.** List of primary antibodies used in the study. Specifying host, isotype, epitope retrieval, staining pattern, source and reference.

| Antibody | Host | Isotype | Epitope retrieval | Staining pattern | Source | Reference |
| --- | --- | --- | --- | --- | --- | --- |
| Anti-DCX | Rabbit | Polyclonal | DCX, neuroblast marker | Cytoplasmic | Abcam (Cambridge, UK) | ab18723 |
| Anti-S100β | Rabbit | Polyclonal | S100β, astrocyte  marker | Cytoplasmic | Abcam (Cambridge, UK) | ab41548 |
| Anti-NeuN | Rabbit | Monoclonal | NeuN, neuronal marker | Nuclear | Abcam (Cambridge, UK) | ab177487 |
| Anti-SOX2 | Rabbit | Polyclonal | SOX2,  astrocyte/stem cell marker | Nuclear | Santa Cruz Biotechnology, Santa Cruz, CA, USA) | sc-20088 |
| Anti-SOX2 | Goat | Polyclonal |  | Nuclear | RD Systems Minneapolis, CO, USA) | AF2018 |
| Anti-MCM2 | Mouse | Monoclonal | MCM2, cell proliferation marker | Nuclear | BD Bioscience  ( East Rutherford, NJ, USA) | 610701 |
| Anti-GFAP | Chicken | Polyclonal | GFAP, glial marker | Cytoplasmic | Abcam (Cambridge, UK) | ab4674 |
| Anti-NeuN | Rat | Monoclonal | NeuN, neuronal marker | Nuclear | Abcam (Cambridge, UK) | ab279297 |
| Anti-BrdU | Rat | Monoclonal | BrdU, cell proliferation marker | Nuclear | Abcam (Cambridge, UK) | ab6362 |

**Supplementary table 2: Secondary antibodies.** List of secondary antibodies used in the study. Specifying host, dilution used, fluorescence conjugated, source and reference

| Antibody | Host | Dilution | Fluorescence | Source | Reference |
| --- | --- | --- | --- | --- | --- |
| Alexa Flour anti-rabbit | Donkey | 1:1000 | 647 | Invitrogen (Carlsbad, CA, USA) | A-32795 |
| Alexa Flour anti-rabbit | Donkey | 1:1000 | 594 | Invitrogen (Carlsbad, CA, USA) | A-21207 |
| Alexa Flour anti-rabbit | Donkey | 1:1000 | 488 | Invitrogen (Carlsbad, CA, USA) | A-21206 |
| Alexa Flour anti-goat | Donkey | 1:1000 | 647 | Invitrogen (Carlsbad, CA, USA) | A-11058 |
| Alexa Flour anti-mouse | Donkey | 1:1000 | 488 | Invitrogen (Carlsbad, CA, USA) | A-21206 |
| Alexa Flour anti-chicken | Goat | 1:1000 | 647 | Abcam (Cambridge, UK) | ab150175 |
| Alexa Flour anti-rat | Donkey | 1:1000 | 488 | Life Technologies (Eugene, OR, USA) | A-21208 |

.

**Supplementary table 3. Statistical comparisons for the acquisition phase of the Morris water maze test.**

| Group | 2-way ANOVA (dayXsex) | Day effect | Sex effect |
| --- | --- | --- | --- |
| **SAMR1** | [F_(3,306)_=0.138, p=0.937] | F=20.55 p<0.001 | F=37.82; p<0.001 |
| **SAMP8** | [F_(3,314)_=0.257, p=0.857] | F=16.61; p<0.001 | F=7.10; p=0.008 |
| **SAMP8-ER** | [F_(3,337)_=0.957, p=0.413] | F=7.69; p<0.001 | F=2.49; F<0.001 |
